# Fat Extract Modulates Calcium Signaling and Protects Against Hyperactive Osteoclastogenesis in Bone Remodeling with Antioxidant Capacity

**DOI:** 10.1101/2022.07.28.501821

**Authors:** Yiqi Yang, Mingming Xu, Tianyou Kan, Yao Wang, Shuhong Zhang, Mingwu Deng, Wenjie Zhang, Hanjun Li, Zhifeng Yu

## Abstract

Osteoclasts are cells which are primarily involved in bone remodeling and osteolytic bone diseases. Hyperactive osteoclastogenesis leads to pathological bone loss and microarchitectural deterioration, particularly in postmenopausal osteoporosis. However, given the limitations of current first-line osteoclast inhibitors, there is an urgent need for a novel antiresorptive agent with higher efficiency and fewer side effects. Cell-free fat extract (CEFFE) is the liquid fraction obtained from human adipose tissues, which are enriched with a variety of cytokines and growth factors. This study aims to explore its pharmaceutical effect on hyperactive osteoclast formation *in vivo* and *in vitro*. CEFFE exhibits excellent potentials to attenuate osteoclast-associated bone loss in an ovariectomy (OVX) mouse model and to inhibit RANKL-induced osteoclastogenesis in primary bone marrow-derived monocytes. Furthermore, the cationic protein fraction of CEFFE (CEFFE-Cation) is identified as the main inhibitory component in osteoclast formation assay. Excessive reactive oxygen species (ROS) production is the main cause of osteoclast overactivation. According to LC-MS/MS analysis, the CEFFE-Cation fraction mainly consists of various antioxidant enzymes, extracellular matrix, and secreted cytokines, which endow it with a superior antioxidant capacity. Ca^2+^ signaling contributes to osteoclast maturation. In addition to scavenging ROS, CEFFE-Cation is also capable of mitigating Ca^2+^ oscillation, calcineurin activation, and subsequent NFATc1 nuclear translocation during osteoclastogenesis. Overall, this study elucidates the promising translational potential of CEFFE as a next-generation personalized antiresorptive agent for osteolytic bone disease treatment.

## 1. Introduction

Hyperactive osteoclastogenesis causes unbalanced bone remodeling, which is a common feature of osteolytic bone diseases, such as osteoporosis, osteoarthritis, tumorous bone metastasis, and osteomyelitis [1-3]. Osteoporosis is a highly prevalent metabolic bone disease characterized by bone loss and deterioration of bone microarchitecture. Globally, an estimated 200 million women have been diagnosed with osteoporosis; one in three women over the age of 50 years suffers from osteoporosis-related fractures, as do one in five men [4]. Osteoporosis is extremely painful and a financial burden to society. Although inhibiting bone resorption is a promising strategy to treat osteoporosis, commercially available antiresorptive medicines are still limited [5]. Presently, denosumab (a recombinant antibody targeting the receptor activator of nuclear factor-κB ligand [RANKL]) and zoledronic acid (a bisphosphonate) are the primary osteoclast inhibitors widely used in the clinic; however, long-term usage of these medicines could lead to multiple side-effects, including mandibular bone necrosis, hypocalcemia, and cystitis [6]. Therefore, the development of a novel antiresorptive agent with higher efficiency and fewer side-effects is urgently needed.

Binding of RANKL to the receptor activator of nuclear factor-κB (RANK) is an essential step in activating the downstream signaling cascades needed for osteoclastogenesis, particularly the accumulation of intracellular reactive oxygen species (ROS) [7]. ROS, as major second messengers, further augment differentiation signals and activate the nuclear factor-κB (NF-κB) and mitogen-activated protein kinase (MAPK) pathways, which crucially participate in osteoclast formation and bone resorption, as well as nuclear factor of activated T-cells 1 (NFATc1) nuclear translocation, driving a wide range of osteoclast-specific gene expression [8]. In recent years, increasing evidence has showed the involvement of other signaling pathways in regulating osteoclast formation and function, among which the research on calcium (Ca^2+^) signaling has attracted the most extensive attention.

Osteoclasts have been considered electrically stable cells, with the Ca^2+^ levels in osteoclasts or their precursors, bone marrow-derived monocytes (BMMs), remaining essentially stable. However, recent studies have found that sequential regenerative changes in intracellular Ca^2+^ levels, also known as Ca^2+^ oscillations, play an indispensable role in regulating osteoclastogenesis [9, 10]. Ca^2+^ oscillation activates calcineurin, a Ca^2+^-dependent serine/threonine phosphatase, which leads to the dephosphorylation of NFATc1. In addition, elevated Ca^2+^ levels are associated with increased intracellular ROS accumulation through mitochondrial impairment and activation of oxidant enzymes [11]. Therefore, targeting Ca^2+^ signaling is a potential therapeutic strategy to prevent hyperactive osteoclastogenesis.

Adipose tissue and bones are closely connected and functionally interdependent in skeletal homeostasis. Adipose-derived factors play a crucial role in orchestrating the balance in bone remodeling [12]. Cell-free fat extract (CEFFE) is a water-soluble liquid mixture containing various growth factors and enzymes obtained from fresh adipose tissue. Owing to its easy accessibility, high biocompatibility, and low immunogenicity, CEFFE holds great promise for improving the treatment of various diseases. Our previous study reported that CEFFE attenuates tail suspension-related bone loss by inhibiting osteocyte apoptosis [13]. In addition, numerous studies have demonstrated that CEFFE can improve graft survival, accelerate diabetic wound healing [14, 15], increase skin thickness [16], enhance tissue regeneration [17], and promote neovascularization [18]. However, CEFFE’s therapeutic effectiveness on osteoclastogenesis hyperactivity remains poorly understood.

In this study, we systemically investigated the pharmaceutical effect and underlying mechanism of CEFFE in hyperactive osteoclast formation and osteolytic bone disease. An ovariectomy (OVX) mouse model was established, which showed that CEFFE efficiently ameliorated osteoclast-associated bone loss and bone microarchitecture deterioration. Mechanistically, CEFFE not only scavenged both extra- and intracellular ROS but also suppressed RANKL-induced Ca^2+^ oscillation, calcineurin activation, and NFATc1 nuclear translocation, resulting in the suppression of hyperactive osteoclastogenesis. This study offers a powerful foundation for future translational applications of CEFFE as next-generation personalized medicine in clinical practice.

## 2. Results

### 2.1. CEFFE protects against bone loss and microarchitecture deterioration in OVX mice

CEFFE was prepared as previously described (**Figure 1A**) [15]. After a series of washes, mechanical emulsification, and centrifugation, fresh fat tissue was converted to water-soluble, non-immunogenic CEFFE. Previous studies confirmed that hyperactive osteoclastogenesis is the leading cause of bone loss in OVX mice [1]. Therefore, to investigate the effect of CEFFE on osteoclastogenesis *in vivo*, OVX mouse models were established and treated with CEFFE twice per week (**Figure 1B**). After six weeks of treatment, micro-CT (μCT) analysis of both tibial and vertebral trabeculae revealed that CEFFE significantly protected against bone loss and trabecular bone deterioration in OVX mice, as evidenced by an increase in the bone volume-to-total volume ratio (BV/TV), connection density (Conn. Dens.), and trabecular number (Tb. N), but a decreased trabecular separation (Tb. Sp) (**Figure 1C-F**). Consistently, μCT analysis of the middle tibia showed a higher cortical thickness (Ct. Th) in the OVX + CEFFE group compared to the OVX + PBS control group (**Figure 1G-H**). Furthermore, not only bone mass and microarchitecture but also bone mechanical properties were enhanced by CEFFE treatment. In a three-point bending assay, femurs from OVX + CEFFE mice exhibited a higher maximal load and elastic modulus than those from OVX + PBS controls (**Figure 1I**). Taken together, these results show that CEFFE efficiently prevented bone loss, microarchitecture deterioration, and bone fragility in OVX mice.

**Figure 1.**
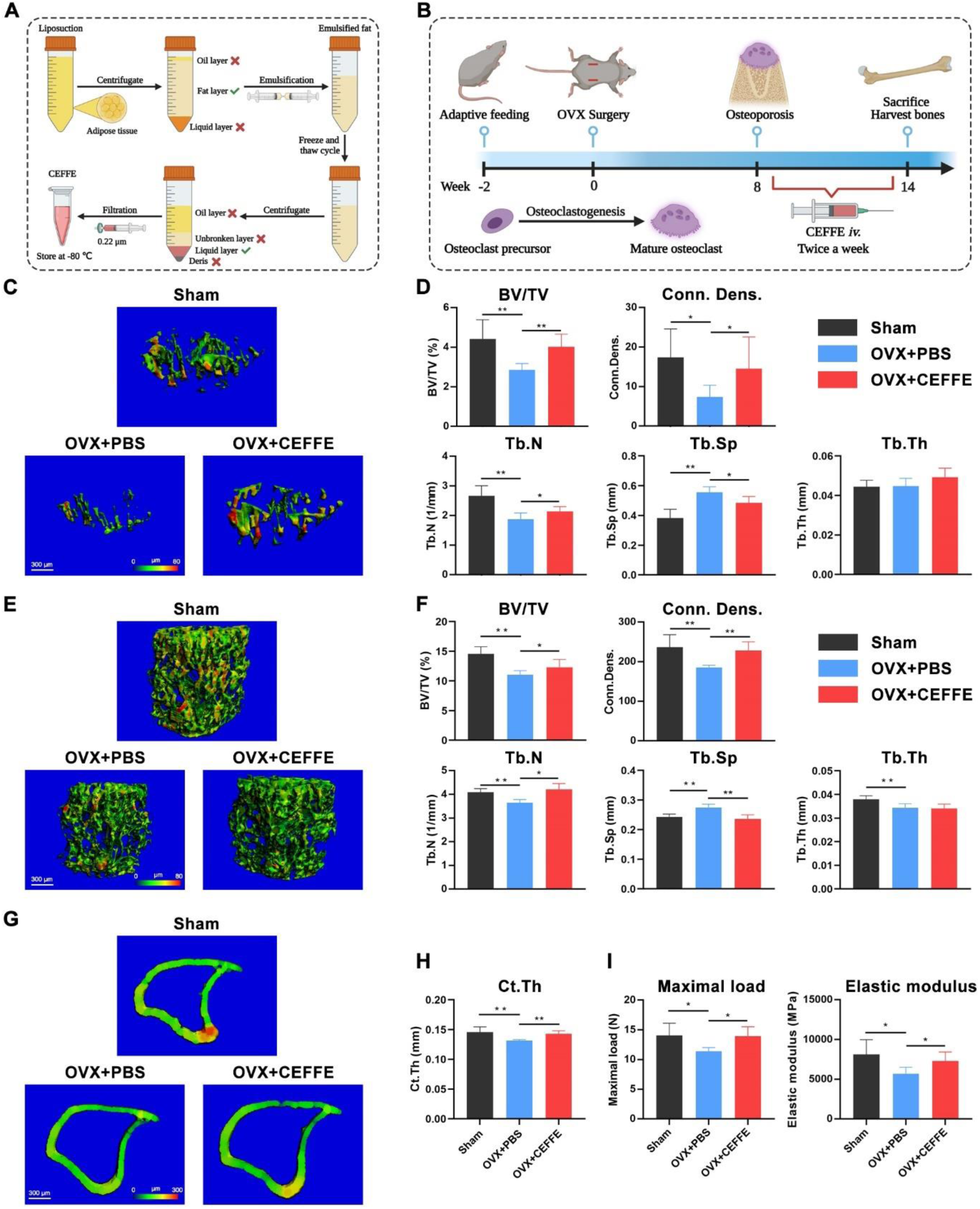
CEFFE protects against bone loss and microarchitecture deterioration in OVX mice. (A) Workflow diagram of CEFFE preparation. (B) Schematic illustration of the animal experiment design. Eight weeks after OVX modeling, the mice were treated with PBS or CEFFE twice a week separately. (C-F) 3D μCT images of tibial trabecular bones and vertebral bodies at six weeks after PBS or CEFFE treatment. Quantitative analysis of bone microarchitecture parameters of trabecular bones and vertebral bodies. (G, H) 3D μCT images of cortical bones from the midshaft of tibia at six weeks after PBS or CEFFE treatment. Quantification of cortical thickness. (I) Maximal load and elastic modulus of femur detected by three-point bending assay at six weeks after PBS or CEFFE treatment. n=8, “*” indicates P < 0.05; “**” indicates P < 0.01.

### 2.2. CEFFE inhibits osteoclastogenesis in OVX mice

Consistent with the μCT results, bone histomorphometric analysis according to hematoxylin & eosin (H&E) staining also revealed that BV/TV was significantly elevated after CEFFE treatment in OVX mice (**Figure 2A, B**). Simultaeously, amelioration of bone marrow adipose tissue enrichment by CEFFE treatment was also observed. To confirm the inhibitory effect of CEFFE on osteoclast formation, we examined osteoclasts on trabecular bone and periosteal bone using tartrate-resistant acid phosphatase (TRAP) staining (**Figure 2C**). As shown in **Figure 2D**, CEFFE treatment markedly decreased the TRAP^+^ osteoclast number in the OVX mice. Dendritic cell-specific transmembrane protein (DC-STAMP) is an essential inducer and core regulator of fused polykaryon formation during osteoclastogenesis [19]. To further validate our observations, we performed immunofluorescence (IF) staining of DC-STAMP in tibial sections (**Figure 2E)**. As expected, OVX-induced DC-SATMP expression was attenuated by CEFFE treatment (**Figure 2F**). In summary, these data indicate that CEFFE markedly inhibited osteoclastogenesis in OVX mice.

**Figure 2.**
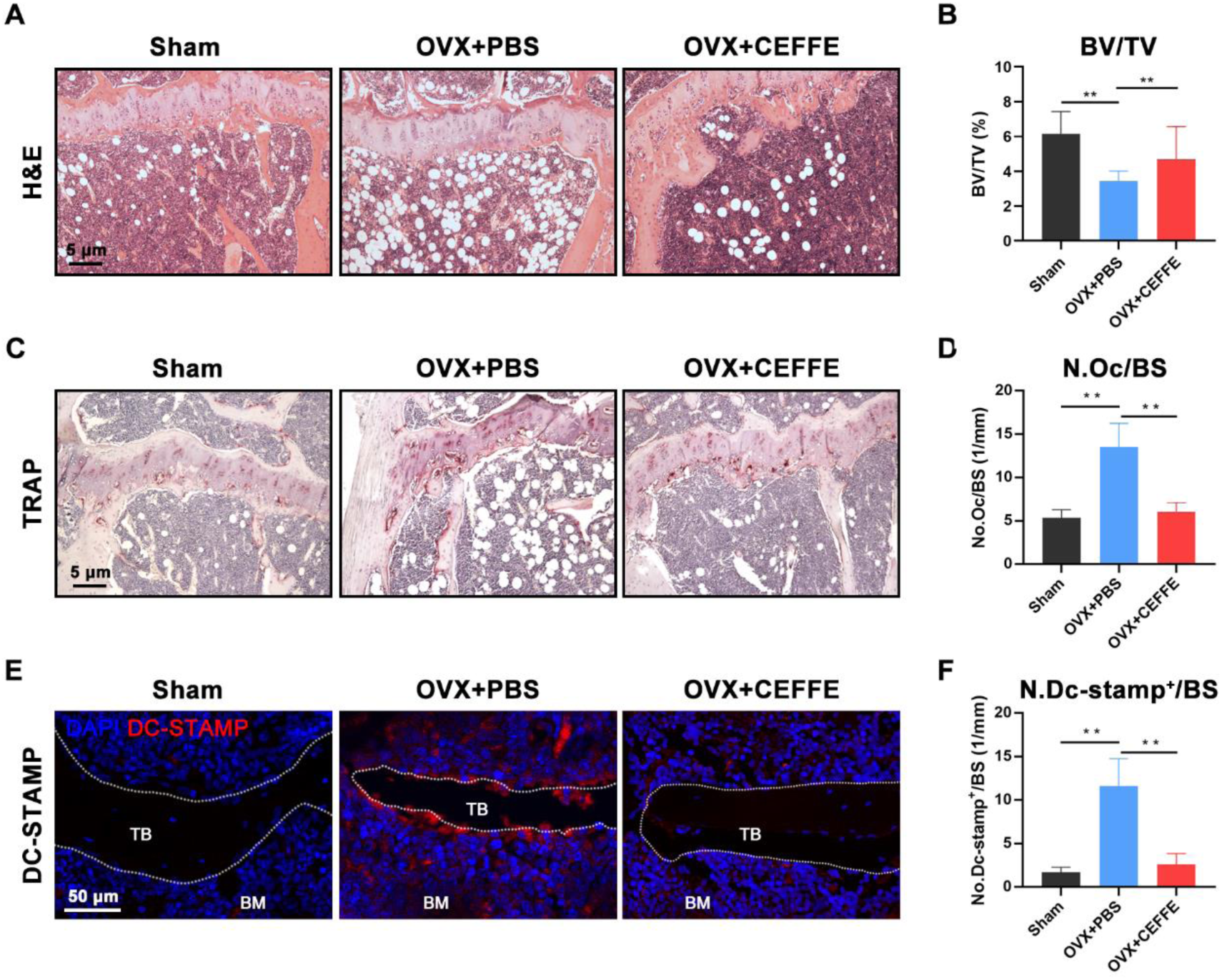
CEFFE inhibits osteoclastogenesis in OVX mice. (A, B) H&E staining of tibial sections. Quantification of trabecular BV/TV based on H&E staining. (C, D) TRAP staining of tibial sections. The TRAP+ osteoclast/bone surface (N.Oc/BS) was quantified based on TRAP staining. (E, F) IF staining of DC-STAMP on the trabecular bone and periosteal bone. Quantification of DC-STAMP expression based on IF staining. n=8, “**” indicates P < 0.01.

### 2.3. CEFFE suppresses osteoclast formation and bone resorption *in vitro*

To verify our results *in vitro*, we studied the effect of CEFFE on RANKL-induced osteoclastogenesis in primary BMMs. First, the effect of CEFFE on BMM viability was assessed using a cell counting kit-8 (CCK-8) assay. CEFFE below 500 μg/mL exhibited no toxicity to primary BMMs, and no proliferation promotion effect was detected (**Figure S1**, Supporting Information). Next, we treated BMMs with various concentrations of CEFFE during RANKL-induced osteoclastogenesis. As illustrated by TRAP and F-actin ring staining, CEFFE effectively suppressed TRAP^+^ osteoclast differentiation and F-actin ring formation in a dose-dependent manner (**Figure 3A, B, D**). Notably, CEFFE also attenuated osteoclast function *in vitro*, as shown by the decreased bone resorption pit area on hydroxyapatite-coated plates after CEFFE treatment (**Figure 3C, D**). Our previous study reported a wide range of osteoclast-specific genes involved in osteoclastogenesis, including *Trap, Ctsk, Dcstamp, Nfatc1, Atp6a3*, and *Atp6d2* [2, 20]. Their expression increased in a time-dependent manner after induction, indicating a progression of differentiation. Here, osteoclast-specific gene expression was quantified by real time qPCR (RT-qPCR) in BMMs treated with or without CEFFE under RANKL stimulation. As shown in **Figure 3E**, CEFFE significantly decreased the RANKL-induced mRNA expression of these genes in a dose-dependent manner. Our results confirmed the inhibitory effect of CEFFE on osteoclast formation and bone resorption *in vitro*.

**Figure 3.**
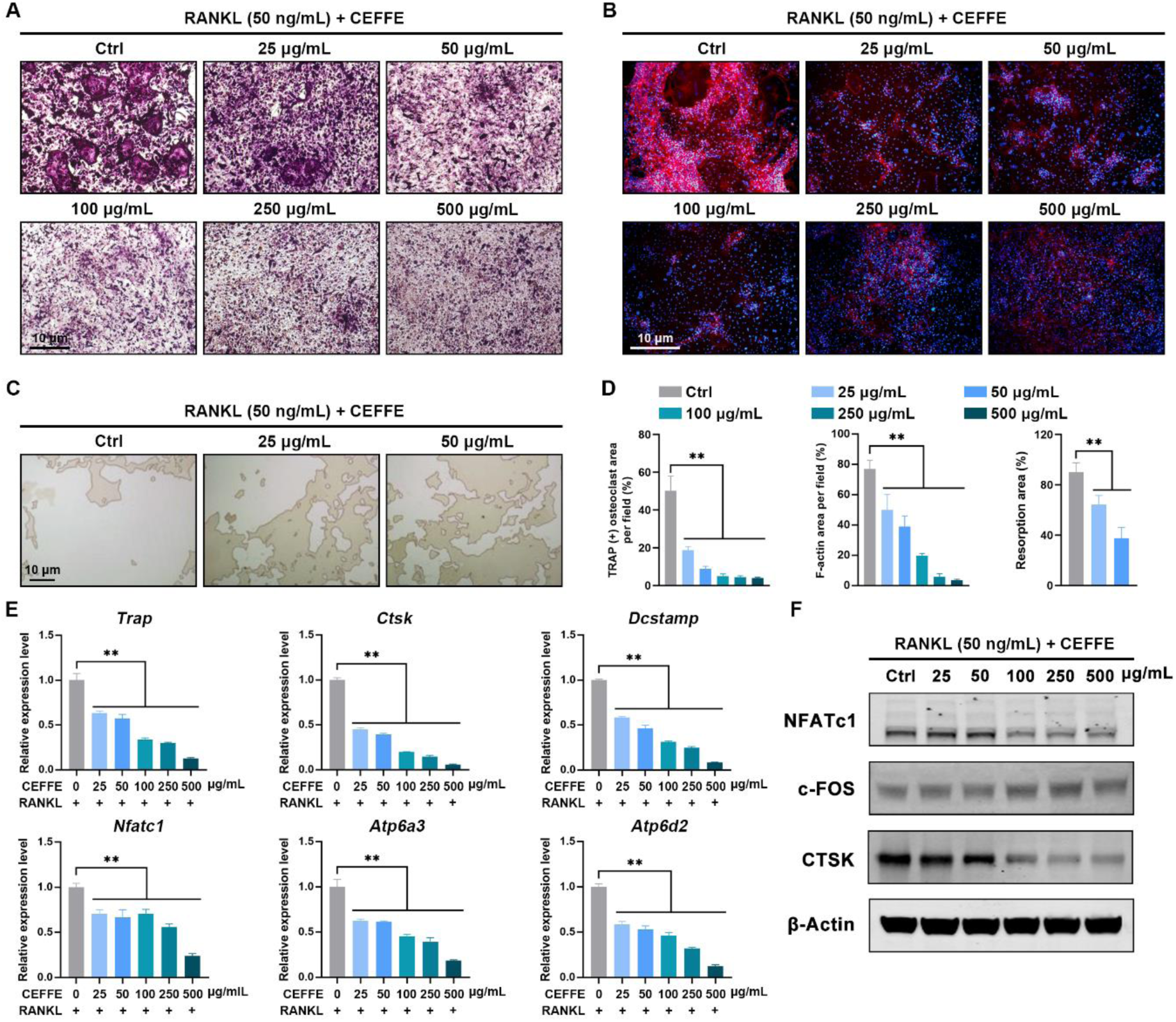
CEFFE suppresses osteoclast formation and bone resorption *in vitro*. (A, B) TRAP staining and F-actin ring staining of BMMs treated with various concentrations of CEFFE under RANKL stimulation. (C) Representative images of bone resorption pits on the hydroxyapatite plates after treatment with various concentrations of CEFFE. (D) Quantification of TRAP^+^ area, F-actin area, and bone resorption area per field. (E, F) Osteoclast-specific genes (*Trap, Ctsk, Dcstamp, Nfatc1, Atp6a3*, and *Atp6d2*) and proteins (NFATc1, c-Fos, and CTSK) expression in BMMs treated with various concentrations of CEFFE under RANKL stimulation. β-Actin was used as the internal control. n=3, “**” indicates P < 0.01.

### 2.4. CEFFE barely affects RANKL-induced MAPK or NF-κB activation

NFATc1 and c-FOS are two major transcriptional factors driving osteoclast-specific gene expression; capthesin K, also known as CTSK, is responsible for the digestion of the bone matrix during bone resorption [21]. As shown by immunoblotting, CEFFE suppressed NFATc1 and CTSK protein expression, which was consistent with the RT-qPCR results. Surprisingly, c-FOS protein expression was almost unaffected by CEFFE treatment (**Figure 3F**). Extracellular signal-related kinase (ERK), a major kinase in MAPK signaling, is a direct upstream kinase that drives c-FOS nuclear translocation. Next, we investigated whether CEFFE influenced RANKL-induced ERK activation (**Figure S2**, Supporting Information). Again, we observed that CEFFE had little effect on ERK phosphorylation. This indicated that the pharmaceutical effect of CEFFE may not be dependent on the ERK/c-FOS axis.

RANKL-dependent NF-κB activation is another major signalling pathway that initiates osteoclastogenesis [22]. This finding was unexpected and suggested that the NF-κB pathway activation was also unaffected by the presence of CEFFE (**Figure S3**, Supporting Information). In addition to MAPK/NF-κB signaling, AKT phosphorylation has also been reported to be an essential activator of osteoclast formation [23]. Although downstream NFATc1 expression was decreased after CEFFE treatment, upstream RANKL-induced AKT activation was still unaltered (**Figure 3F**; **Figure S2**, Supporting Information). Collectively, we postulate that the pharmaceutical effect of CEFFE might be mediated by NFATc1 but independent of MAPK, NF-κB, or AKT signaling.

### 2.5. CEFFE-Cation is the major active component for inhibiting osteoclast formation

CEFFE is a mixture of multiple proteins and non-coding RNA. To identify the major bioactive component of CEFFE, we first pre-incubated CEFFE with proteinase K (PK) or RNase A to separately remove the protein or RNA. The pre-digested products were then tested for RANKL-induced osteoclast differentiation. TRAP staining revealed that PK pretreatment efficiently attenuated the inhibitory effect of CEFFE on osteoclast formation, whereas RNase A pretreatment had little or no rescue effect (**Figure 4A**). Similarly, osteoclasts in the PK pretreatment group exhibited higher osteoclast-specific gene expression levels than those in cells incubated with pure CEFFE (**Figure 4B**). This result suggests that the proteins in CEFFE may be the main components responsible for inhibiting osteoclastogenesis.

**Figure 4.**
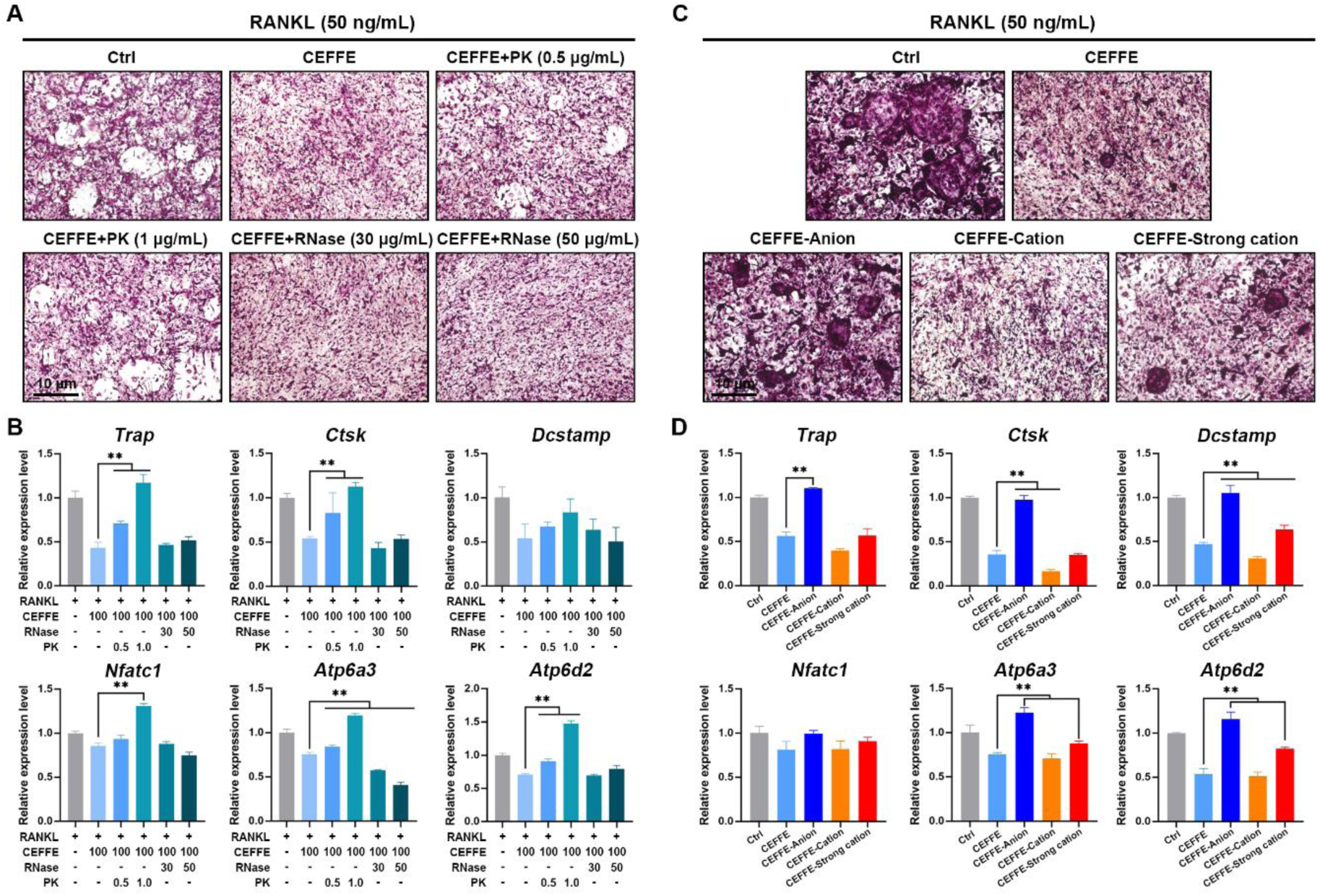
CEFFE-Cation is the major active component for inhibiting osteoclast formation. (A, B) CEFFE was predigested with proteinase K or RNase A to remove protein or RNA components separately. CEFFEs with different components (100 μg/mL) were tested via an osteoclast formation assay under RANKL stimulation. TRAP staining and osteoclast-specific gene (*Trap, Ctsk, Dcstamp, Nfatc1, Atp6a3*, and *Atp6d2*) expression of BMMs were presented. (C, D) CEFFE was further subdivided into anionic proteins (CEFFE-Anion), cationic proteins (CEFFE-Cation), and strong cationic proteins (CEFFE-Strong cation) by ion-exchange chromatography. CEFFEs with different surface charges (100 μg/mL) were tested in osteoclast formation assays under RANKL stimulation. TRAP staining and osteoclast-specific gene (*Trap, Ctsk, Dcstamp, Nfatc1, Atp6a3*, and *Atp6d2*) expression of BMMs were presented. n=3, “**” indicates P < 0.01.

In contrast to RNA, which is negatively charged at physiological pH, the surface charge of a protein is determined by its isoelectric point. Based on the surface charge, CEFFE was subdivided into its anionic components (CEFFE-Anion), cationic components (CEFFE-Cation), and strong cationic components (CEFFE-Strong cation) using ion-exchange chromatography. Furthermore, CEFFE components with different surface charges were tested for RANKL-induced osteoclast differentiation. As demonstrated in **Figure 4C**, the minimal TRAP^+^ osteoclast area was observed in the CEFFE-Cation group, followed by the CEFFE-Strong cation group, while CEFFE-Anions showed no inhibitory effect on osteoclast formation. Simultaneously, the same trend was observed in osteoclast-specific gene expression as detected by RT-qPCR (**Figure 4D**). Together, these data suggest that the CEFFE-Cation group, which mainly consists of positively charged proteins, is the major active component that inhibits osteoclast formation.

### 2.6. Mass spectrometric identification of proteins in CEFFE-Cation

To investigate the detailed mechanism of the CEFFE-Cation group in inhibiting osteoclastogenesis, we performed Coomassie brilliant blue staining to visualize the bands of separated proteins, followed by liquid chromatography with tandem mass spectrometry (LC-MS/MS) analysis. The proteins were enriched below 95 kD, indicating that the CEFFE-Cation group was dominated by small-molecule proteins. Notably, the mass spectrum analysis identified more than 500 proteins in the CEFFE-Cation group, and the highly enriched representative proteins are listed in **Figure 5A**. Most of the identified proteins were antioxidant enzymes, extracellular matrix proteins, and bioactive secreted cytokines. Meanwhile, several molecular functions and biological processes associated with antioxidation were significantly enriched in GO analysis, whereas KEGG analysis demonstrated that pathways related to cell metabolism and survival were preferentially targeted (**Figure 5B, C**). ROS in the microenvironment are essential factors that promote osteoclastogenesis. Therefore, we speculated that CEFFE-Cations functions by scavenging ROS to inhibit osteoclast formation. To validate our hypothesis, we assessed the total antioxidant capacity of CEFFE-Cations in a cell-free system. The results suggested that CEFFE-Cations displayed a superior antioxidant ability in a dose-dependent manner (**Figure 5D**).

**Figure 5.**
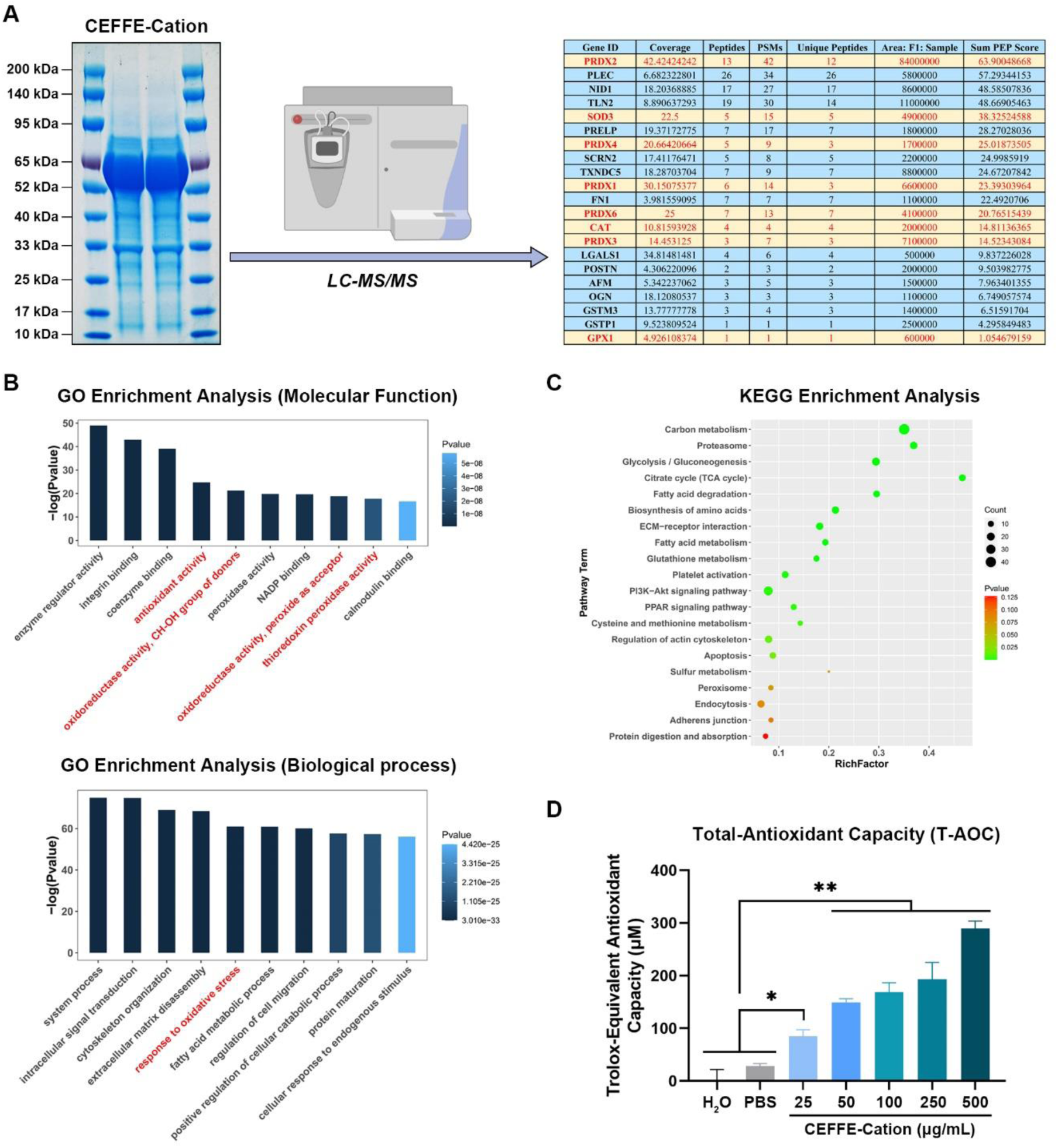
Mass spectrometric identification of proteins in the CEFFE-Cation fraction. (A) Coomassie brilliant blue staining of fractionated CEFFE-Cations was presented. Furthermore, 21 specified functional proteins in the CEFFE-Cation fraction were identified by LC-MS/MS analysis. (B, C) GO and KEGG enrichment analysis based on the identified proteins in the CEFFE-Cation group by LC-MS/MS. (D) The total-antioxidant capacity of the CEFFE-Cation group was assessed in a cell-free system. n=3, “*” indicates P < 0.05; “**” indicates P < 0.01.

### 2.7. CEFFE-Cation mitigates intracellular ROS accumulation and NFATc1 nuclear translocation during RANKL-induced osteoclastogenesis

To further study the antioxidant ability of CEFFE-Cations in the cell system, intracellular ROS accumulation was assessed using a dichlorodihydrofluorescein diacetate (DCFH-DA) probe in BMMs treated with and without CEFFE-Cations during RANKL-induced osteoclastogenesis. As expected, RANKL stimulation significantly augmented intracellular ROS production, whereas CEFFE-Cations alleviated ROS accumulation in BMMs in a time-dependent manner (**Figure 6A**). Consistently, immunoblotting revealed that CEFFE-Cations markedly promoted the expression of major antioxidant enzymes, including SOD1, CAT, and GPX4 (**Figure 6B**). This indicates that CEFFE-Cation not only scavenges ROS in the microenvironment but also enhances the antioxidant capacity of BMMs during RANKL-dependent osteoclastogenesis.

**Figure 6.**
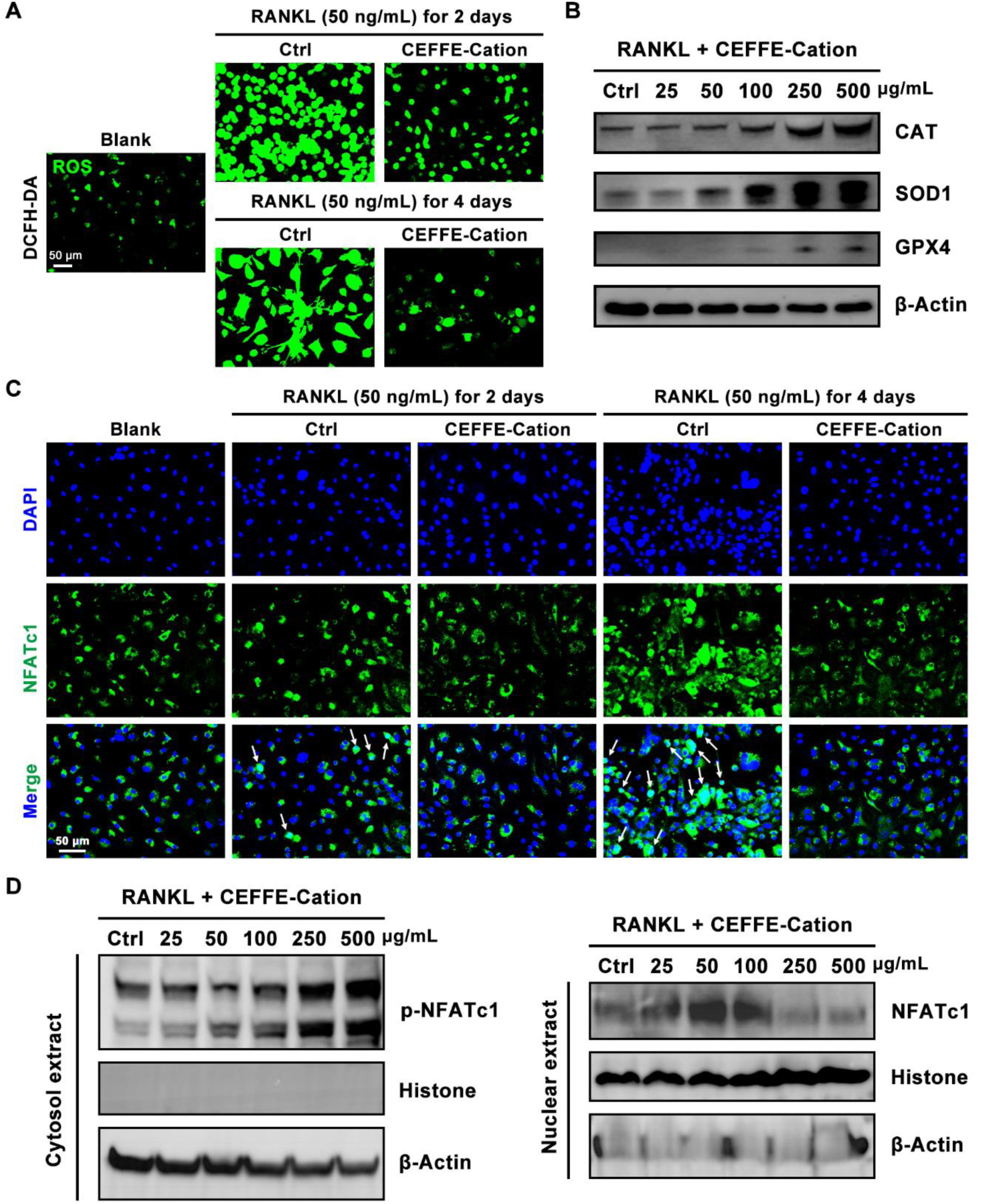
CEFFE-Cations mitigate intracellular ROS accumulation and NFATc1 nuclear translocation during RANKL-dependent osteoclastogenesis. (A) Fluorescent ROS staining of BMMs treated with/without CEFFE-Cations after two or four days’ RANKL stimulation. (B) Expression of major antioxidant enzymes (CAT, SOD1, and GPX4) in BMMs after CEFFE treatment. (C) Immunofluorescent staining of NFATc1 in BMMs treated with and without CEFFE-Cations after two or four days’ RANKL stimulation. (D) Immunoblotting showing p-NFATc1 expression in cytosol and NFATc1 enrichment in nucleus after CEFFE treatment. n=3.

ROS has been reported to augment NFATc1 activation [24]. As shown in **Figure 3F**, we observed a decrease in NFATc1 expression in BMMs after CEFFE treatment. However, whether CEFFE-Cations attenuate RANKL-induced NFATc1 activation remains unclear. Here, we examined the effect of CEFFE-Cations on NFATc1 nuclear translocation during RANKL-dependent osteoclastogenesis by detecting the colocalization of DAPI (blue fluorescence) and the NFATc1 signal (green fluorescence). As shown in **Figure 6C**, RANKL-induced nuclear import of NFATc1 was significantly diminished by CEFFE-Cation treatment. Dephosphorylation of NFATc1 results in a conformational change that exposes its nuclear localization signal and promotes nuclear import. To further verify our results, the nuclear and cytoplasmic proteins of BMMs treated with and without CEFFE-Cations under RANKL stimulation were extracted for immunoblotting analysis. As expected, CEFFE-Cations increased the phosphorylation of NFATc1 at Ser294 and inhibited its nuclear translocation (**Figure 6D**). The above experimental data indicate that CEFFE-Cations simultaneously mitigate intracellular ROS accumulation and NFATc1 nuclear translocation during RANKL-dependent osteoclastogenesis.

### 2.8. CEFFE-Cation suppresses Ca2+ oscillation and calcineurin activity during RANKL-induced osteoclastogenesis

In addition to ROS, calcium signaling is another major upstream pathway promoting NFATc1 nuclear import and RANKL-induced osteoclastogenesis [25]. To investigate whether CEFFE attenuated RANKL-induced NFATc1 activation by modulating Ca^2+^signaling, we first assessed intracellular Ca^2+^ levels using a Fluo-4AM probe in BMMs treated with and without CEFFE-Cations during RANKL-dependent osteoclastogenesis. RANKL stimulation induced a sustained elevation of Ca^2+^, while CEFFE-Cations reversed this trend and caused a reduction in intracellular Ca^2+^ (**Figure 7A, C**). This experiment was conducted from a static perspective. Next, we analyzed RANKL-induced Ca^2+^ oscillations in BMMs from a dynamic perspective. Intracellular Ca^2+^ changes were continuously monitored within 8 min in the presence or absence of CEFFE-Cations. As demonstrated in **Figure 7B**, RANKL evoked Ca^2+^ oscillations, whereas CEFFE-Cations completely abolished this RANKL-induced Ca^2+^ oscillation (**Figure 7D**; **Movie S1-S3**, Supporting Information). Generally, Ca^2+^ is a direct signaling molecule that activates calcineurin activity, and NFATc1 is regulated through Ca^2+^ or calcineurin-dependent serine dephosphorylation. As detected by RT-qPCR, CEFFE-Cations decreased the expression of calcium signaling-related genes, including calmodulin mRNA (*Calm1*) and calcineurin mRNA (*Rcan1, Caln1*, and *Cnb1*) (**Figure 7E**). Notably, CEFFE-Cation also exhibited a direct inhibitory effect on calcineurin activity in BMMs following RANKL stimulation (**Figure 7F**). In summary, these findings imply that CEFFE not only scavenges both extra- and intracellular ROS but also suppresses RANKL-dependent Ca^2+^ oscillations, calcineurin activation, and NFATc1 nuclear translocation, resulting in efficient inhibition of osteoclastogenesis (**Figure 8**).

**Figure 7.**
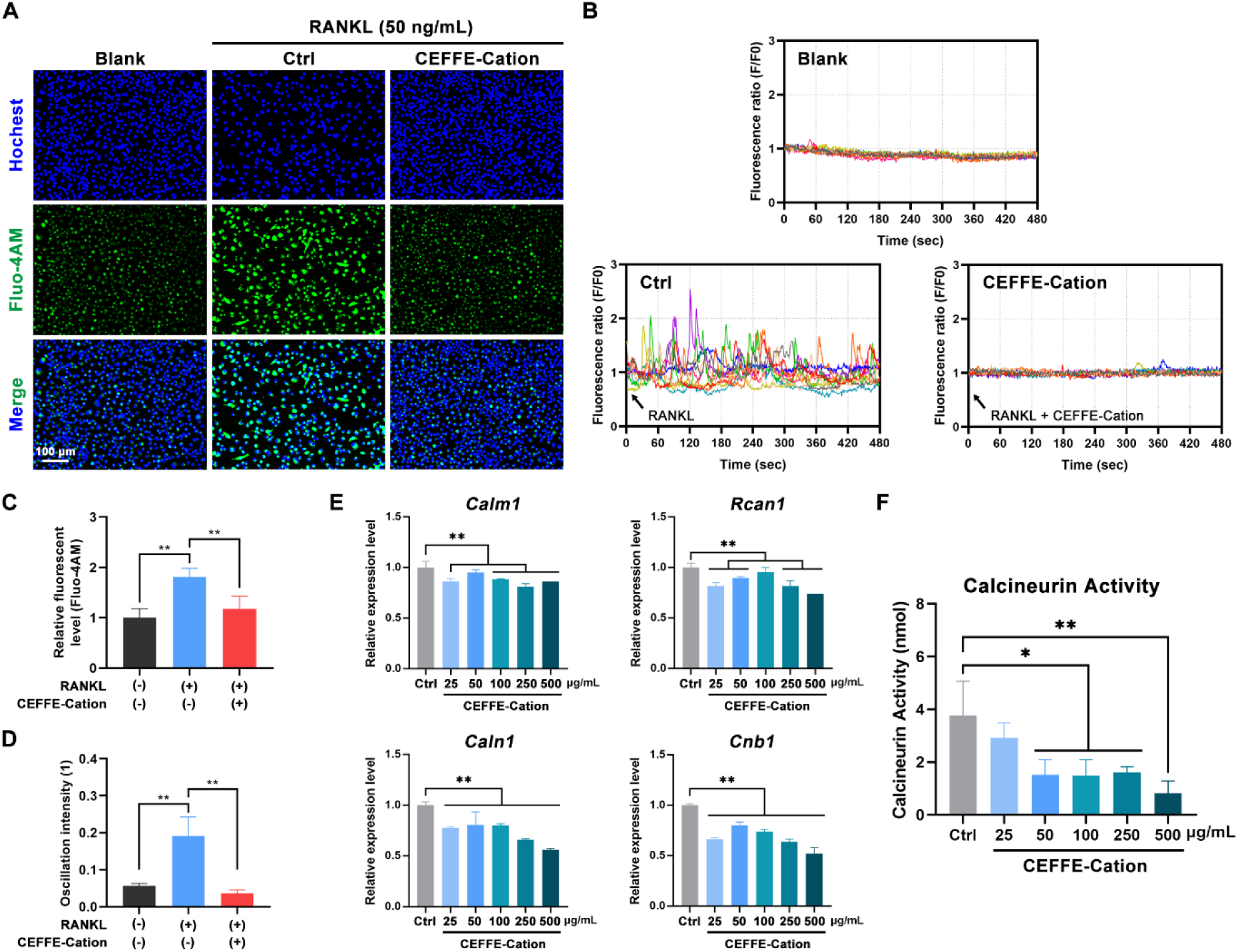
CEFFE-Cations suppress Ca^2+^ oscillation and calcineurin activity during RANKL-dependent osteoclastogenesis. (A) Fluorescent staining of Ca^2+^ in BMMs treated with and without CEFFE-Cations under RANKL stimulation. (B) BMMs under the indicated treatments were loaded with Fluo-4AM fluorescent probe for Ca^2+^ imaging. The intracellular Ca^2+^ signals were continuously monitored for eight min. Each color indicates an individual cell in the field. Notably, Ca^2+^ oscillation evoked by RANKL was significantly inhibited by CEFFE-Cations. (C) Quantification of intracellular Ca^2+^ levels based on (A). (D) Quantification of Ca^2+^ oscillation intensity in the field based on (B). (E) The expression of calcium signaling-related genes (*Calm1, Rcan1, Caln1*, and *Cnb1*) was detected by RT-qPCR after CEFFE-Cation treatment. (F) The assessment of calcineurin activity in BMMs treated with various concentrations of CEFFE-Cation during RANKL-dependent osteoclastogenesis. n=3, “*” indicates P < 0.05; “**” indicates P < 0.01.

**Figure 8.**
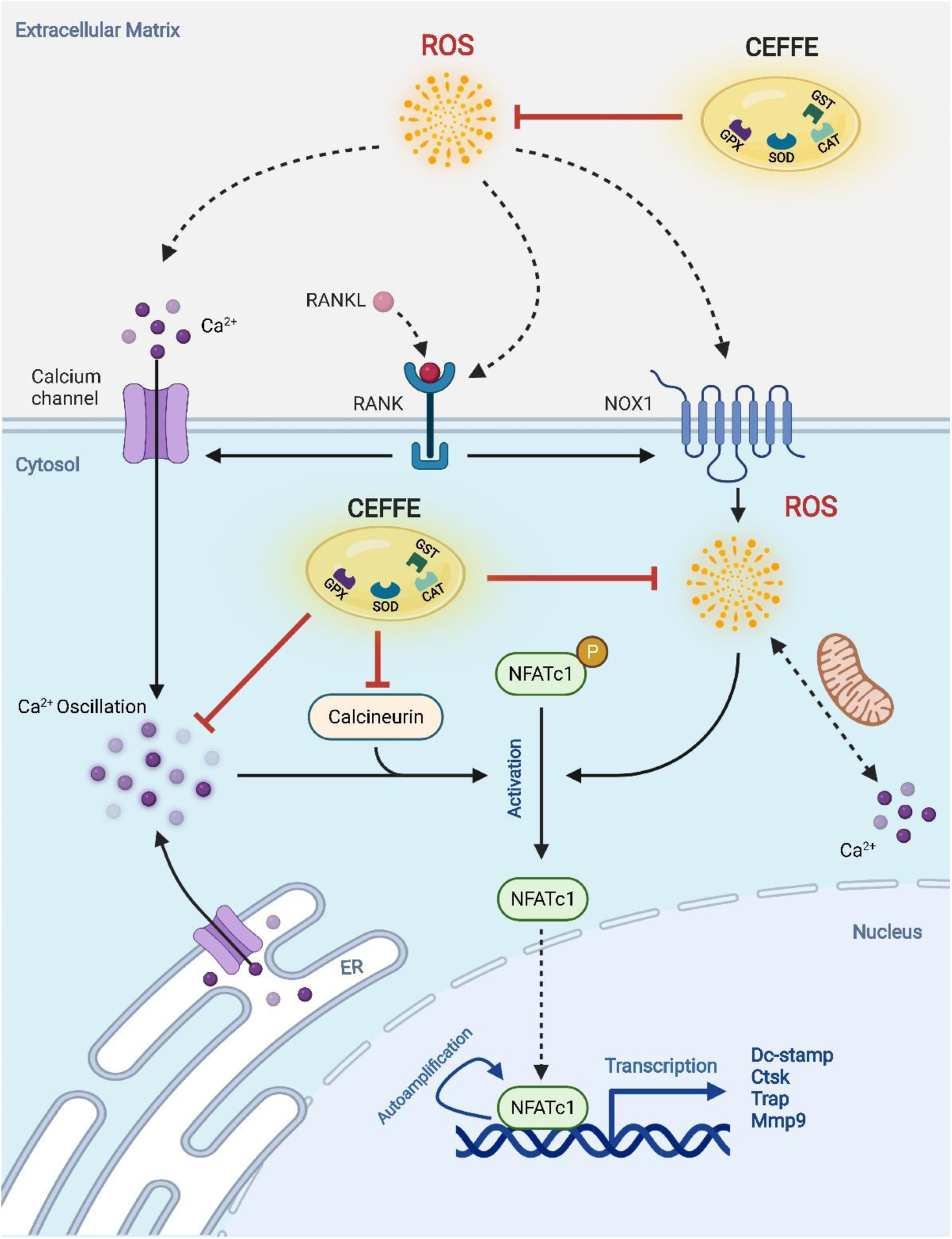
Schematic diagram showing the mechanism by which CEFFE protects against hyperactive osteoclastogenesis and modulates calcium signaling with antioxidant capacity. As identified by mass spectrometry, CEFFE contains a wide range of antioxidant enzymes and secreted cytokines. Mechanistically, CEFFE not only scavenges both extra- and intracellular ROS, but also suppresses RANKL-dependent Ca^2+^ oscillation, calcineurin activation and NFATc1 nuclear translocation, resulting in the efficient inhibition of osteoclastogenesis.

## 3. Discussion

Since the turn of the century, we have witnessed increasing evidence illustrating the crosstalk between fat and bone. Counterintuitive to what had been previously understood, numerous studies have reported the essential role of adipose tissue in maintaining bone homeostasis in various diseases [26, 27]. Thus, recent efforts have focused on developing new therapeutics based on fat extracts to modulate imbalanced bone remodeling. In this study, we identified positively charged proteins as the major bioactive component of cell-free fat extract (CEFFE), and further provided a direct insight into its pharmaceutical effect on inhibiting osteoclast formation. CEFFE treatment protects against bone loss, bone microarchitecture deterioration, and hyperactive osteoclastogenesis in OVX mice. Mechanistically, CEFFE, with its superior antioxidant capacity, effectively suppressed RANKL-induced ROS accumulation, and calcium signaling activation, resulted in the inhibition of NFATc1-dependent osteoclast differentiation.

Adipose tissue is not only a fat-storage organ but also an endocrine organ that produces adipocytokines and metabolites. However, the beneficial role of fat products remains debatable. Obesity severely impairs skeletal health and serum lipid levels are inversely correlated with bone mass [28]. In contrast, a study by Kajimura *et al*. reported that adipose-derived adiponectin positively affected bone health at a young age with little effect at an older age [29]; Upadhyay *et al*. found that adipose-derived leptin provided a therapeutic option for osteoporosis in both human and animal studies [30]; and Kirk *et al*. revealed that adipose-derived insulin-like growth factor-1 (IGF-1) increased bone formation [31]. These different observations are not contradictory. We concluded that the effect of adipose tissue on bone homeostasis is highly age-related and dose-dependent; the adverse influence of fat tissue mainly comes from lipid and liposoluble metabolites. In this study, the lipid component of CEFFE was eliminated by serial centrifugation. According to LC-MS/MS analysis, the CEFFE-Cation fraction primarily comprised of water-soluble small-molecule proteins, including antioxidant enzymes, extracellular matrix, and bioactive adipokines. CEFFE preparation procedures not only retained the effective components but also minimized the deleterious influence of fat tissue. As a next step, we plan to elucidate the exact role and mutual effects of different CEFFE components.

Although fat transplantation and adipose-derived stem cells (ADSCs) already exhibit superior therapeutic value for many diseases, their clinical application is hindered by the strong immunogenicity of adipose tissues and cells [32]. Allograft rejection brings a huge risk and healthy burden to patients. In contrast to adipose tissue, CEFFE contains no cells and few large-molecule proteins; therefore, it has been proven to be safe with a wide range of indications and minimal complications. In addition, CEFFE is characterized by easy accessibility, wide range of sources, and simple preparation procedures. These advantages ensure the effectiveness of CEFFE and further enhance its feasibility for clinical translation.

In addition to inhibiting osteoclast formation, CEFFE has also been reported to possess anti-apoptotic, anti-inflammatory, and pro-proliferative effects. Xu *et al*. reported that CEFFE increased dermal thickness by promoting angiogenesis [16]. Xu *et al*. demonstrated that CEFFE prevents osteocyte apoptosis in a tail-suspension mouse model by attenuating ROS production [13]. Yin *et al*. found that CEFFE promoted diabetic ulcer healing by enhancing re-epithelization [14]. In this study, we elucidated the mechanism by which CEFFE suppressed RANKL-induced ROS production and NFATc1 nuclear import. Considering that aging is accompanied by ROS production and cell apoptosis, CEFFE might be an efficient therapeutic agent for degenerative diseases, including osteoarthritis, vertebral disc degeneration, and diabetic osteoporosis. In the future, we intend to investigate the therapeutic effects of CEFFE in various clinical applications.

Interestingly, we noticed that CEFFE barely affected RANKL-induced ERK, AKT, and NF-κB activation. This is acceptable because these signaling pathways are important, but not exclusive factors that initiate osteoclast differentiation. Consistent with our results, Li *et al*. reported that cell-penetrating peptides abrogated RANKL-stimulated NFATc1 expression without affecting RANKL-induced activation of the NF-κB, ERK, or Akt signaling pathways [33]. Similarly, Lee *et al*. found that lumican inhibited osteoclastogenesis but did not affect ERK phosphorylation [34]; Kim *et al*. also demonstrated that melatonin suppressed osteoclast formation in an ERK/c-FOS-independent way [35]. Therefore, to further elucidate the pharmaceutical mechanism of CEFFE, we focused on Ca^2+^ signaling, which was known to initiate osteoclastogenesis independently of NF-κB, ERK and Akt signaling.

Calcium is an essential secondary messenger involved in a wide range of cell processes. Ca^2+^ signaling controls the survival, differentiation, fusion, and activation of osteoclasts [36]. NFATc1 nuclear import is a significant event in osteoclastogenesis [37]. Ca^2+^ oscillation facilitates the dephosphorylation of NFATc1 in a Ca^2+^/calcineurin-dependent way and promotes its nuclear import. In this study, CEFFE-Cations attenuated both intracellular Ca^2+^ accumulation and RANKL-induced Ca^2+^ oscillation in BMMs. Furthermore, calcineurin activity was also inhibited by CEFFE-Cations, resulting in the blocking of NFATc1 nuclear translocation and inhibition of its auto-amplification. Ca^2+^ signaling and ROS share a mutually promoting relationship [38]. The CEFFE-Cation group consists of various antioxidant enzymes, not only scavenges ROS in the microenvironment but also promotes endogenous antioxidant enzyme expression, further enhancing the reserve antioxidant capacity of BMMs. Due to the simultaneous blockage of Ca^2+^ signaling and ROS accumulation, CEFFE exhibited a promising therapeutic effect of inhibiting bone resorption in unbalanced bone remodeling, as evidenced by the efficient protection against bone loss and microarchitecture deterioration in OVX mice. In summary, our study elucidated the pharmaceutical mechanism by which CEFFE prevents hyperactive osteoclastogenesis. Notably, CEFFE successfully protected OVX mice from bone loss and fragility. Mechanistically, CEFFE not only scavenges microenvironmental ROS but also suppresses RANKL-induced Ca^2+^ oscillation, calcineurin activation, and NFATc1 nuclear translocation. Thus, CEFFE holds promising clinical translational prospects as a next-generation personalized medicine for osteolytic disease treatment.

## 4. Experimental Section

### 4.1 CEFFE Preparation

CEFFE was prepared and provided by Shanghai Stem Cell Technology Co., Ltd. (Shanghai, China). Briefly, fresh fat tissue was collected from healthy donors during liposuction procedures with fully informed consent. Then, the fat samples were washed, mechanically emulsified, centrifuged multiple times, and filtered through a 0.22 μm sterilizing filter (Corning, USA). The liquid phase of the fat extract was collected and stored at -80°C. The CEFFE protein concentration was determined using an enhanced BCA protein assay kit (Beyotime, China). To identify the distinct functions of different components of CEFFE, we predigested CEFFE with proteinase K (ST533, Beyotime, China; 0.5 μg/mL or 1.0 μg/mL) or RNase A (ST577, Beyotime, China; 30 μg/mL or 50 μg/mL) to isolate the protein and RNA components separately. Next, to further subdivide the protein components in CEFFE according to their isoelectric points, ion-exchange chromatography was performed using a HiTrap Q HP column (GE Healthcare, USA). In this process, CEFFE was divided into its anionic components (CEFFE-Anion), cationic components (CEFFE-Cation), and strong cationic components (CEFFE-Strong cation). All CEFFEs with different components were used in the subsequent biofunctional study.

### 4.2 Mice and Ovariectomy (OVX) Model

All animal experimental procedures were approved by the Animal Ethics Committee of Shanghai Ninth People’s Hospital (ethical approval number: SH9H-2020-A754-1). Twenty-four 12-week-old C57BL/6J mice were obtained from Shanghai SIPPR BK Laboratory Animals Ltd. (Shanghai, China) and housed under specific pathogen-free (SPF) conditions throughout this study. After two weeks of adaptive feeding, the mice were randomly subdivided into three groups: a sham control group (Sham; n = 8), an OVX model with PBS treatment group (OVX + PBS; n = 8), and an OVX model with CEFFE treatment group (OVX + CEFFE; n = 8). Each mouse underwent either a sham or bilateral ovariectomy. Eight weeks after surgery, the OVX mice were treated with PBS or CEFFE (50 mg/kg) twice a week. Six weeks after the first CEFFE treatment, all mice were euthanized and their bone samples harvested for subsequent evaluation.

### 4.3 Micro-CT (μCT) Analysis

The effect of CEFFE on the bone microarchitecture was evaluated in accordance with our previously published study [13]. Briefly, right tibias and vertebral bodies were harvested for high-resolution μCT analysis (μCT 80, Scanco, Zurich, Switzerland) at a resolution of 10 μm per voxel, 70 kV voltage, and 114 μA electric current. Quantification of microarchitecture parameters, including BV/TV (%), Tb.N (1/mm), Tb.Th (mm), Tb.Sp (mm), Conn. Dens (1), and Ct.Th (mm), were performed using Scanco software.

### 4.4 Mechanical Property Evaluation

The right femurs from the mice were harvested for the mechanical property test. Twenty-four hours before testing, the bone samples were pre-incubated with 0.9% sodium chloride solution at 37°C. A three-point bending test was performed using a material-testing machine (Model 8874, Instron Corp., USA). The bone samples were loaded to failure at a displacement rate of 1 mm/s, and both load (N) and displacement (mm) were recorded throughout this process.

### 4.5 Histology Analysis

The left limbs of the mice were harvested for histological analysis. Briefly, bone samples were fixed using 4% paraformaldehyde, decalcified, embedded in paraffin, and sectioned into 5 μm-thick slides. Hematoxylin and eosin (H&E) staining and tartrate-resistant acid phosphatase (TRAP) staining were performed to evaluate the protective effect of CEFFE on trabecular bone deterioration. IF staining was performed using antibodies against DC-STAMP (1:200, Millipore, USA). Finally, images were obtained using an optical microscope (Leica, USA), and quantitative bone histomorphometric analysis was performed using the Image-Pro Plus software (Media Cybernetics, USA).

### 4.6 Bone-marrow Derived Macrophage (BMM) Isolation and Osteoclastogenesis

Primary BMMs from the femurs and tibias of 4-week-old C57BL/6J mice were isolated as previously described [39]. The BMMs were cultured in α-MEM medium supplemented with 1% penicillin-streptomycin-gentamycin (Beyotime, China), 10% fetal bovine serum (Sigma, USA), and 25 ng/mL M-CSF (R&D Systems, USA). The medium was changed every three days until the BMMs reached 95% confluency. The viability of BMMs treated with CEFFE was quantified using a CCK-8 assay kit (Beyotime, China) following the manufacturer’s instructions. During osteoclastogenesis induction, BMMs were plated into 96-well or 24-well plates (3 × 10^4^ cells per milliliter) in complete α-MEM culture media supplemented with 50 ng/mL RANKL (R&D Systems, USA) and 25 ng/mL M-CSF (PeproTech, USA). The induction culture lasted for seven days, and the medium was refreshed every three days. On day seven, the mature osteoclasts were processed for subsequent analysis.

### 4.7 TRAP Staining

BMMs were plated in 96-well plates and cultured in osteoclastogenesis media with different treatments. After seven days of induction, mature osteoclasts were fixed for TRAP staining *in vitro*. A TRAP kit (Sigma-Aldrich, USA) was used for staining according to the manufacturer’s instructions. TRAP-staining images were obtained using an optical microscope (Leica, USA). TRAP^+^ osteoclast area per field was quantified using Image-Pro Plus software (Media Cybernetics, USA).

### 4.8 F-actin Ring Analysis

BMMs were plated in 24-well plates and cultured in osteoclastogenesis medium with different treatments. After seven days of induction, mature osteoclasts were fixed with 4% paraformaldehyde for 30 min, rinsed with PBS three times, and stained with rhodamine phalloidin (Abcam, UK) for 20 min. The cells were further counterstained with DAPI (Sigma-Aldrich, USA) for 8 min before observation. Fluorescent images of F-actin rings were captured using a fluorescence microscope (Leica, USA), and the F-actin area per field was quantified using Image-Pro Plus software (Media Cybernetics, USA).

### 4.9 Bone Resorption Analysis

BMMs were plated into hydroxyapatite-coated OsteoAssay plates (Corning, USA) at a density of 3 × 10^4^ cells/mL in osteoclastogenesis media with various treatments. After 10 days of induction, the cells were removed using a soft brush. Bone resorption pits on the hydroxyapatite-coated OsteoAssay plates were observed and captured using a phase-contrast inverted optical microscope (Leica, USA). Quantification of the bone resorption area per field was performed using Image-Pro Plus software (Media Cybernetics).

### 4.10 Real-time qPCR Analysis

RT-qPCR was performed as described previously [40]. BMMs were plated in 24-well plates and cultured in osteoclastogenesis media with various treatments. After seven days of induction, the cells were harvested for total RNA isolation using TRIzol reagent (Invitrogen, USA). Reverse transcription PCR was performed using GoScript Super-Mix (Promega, USA), and RT-qPCR was performed using SYBR Green Super-Mix (Bimake, China) on an RT-qPCR machine (Applied Biosystems, USA). *β-actin* was used as the reference gene. All the primer sequences are listed in **Table S1** (Supporting Information).

### 4.11 Western Blot (WB) Analysis

WB was performed as described in our previous study [41]. BMMs were plated in 24-well plates and cultured in osteoclastogenesis media with various treatments. After seven days of induction, cells were harvested for protein lysis using a nuclear and cytoplasmic protein extraction kit (Beyotime, China) with a protease phosphatase inhibitor cocktail (Beyotime, China). Proteins were then separated using SDS-PAGE, transferred to a PVDF membrane, blocked using 5% skim milk, incubated with primary antibodies, and detected using an infrared imaging platform (Odyssey, USA). The following antibodies were used in this study: anti-NFATc1 (1:1000, Santa Cruz Biotechnology, USA), anti-p-NFATc1 (1:1000, Affinity Biosciences, USA), anti-c-FOS (1:1000, Proteintech, China), anti-CTSK (1:1000, Proteintech, China), anti-β-actin (1:1000, Proteintech, China), anti-histone H3 (1:1000, Abcam, USA), anti-GPX4 (1:1000, Santa Cruz Biotechnology, USA), anti-CAT (1:1000, Proteintech, China), and anti-SOD1 (1:1000, Proteintech, China). For Coomassie brilliant blue staining, the protein bands of the CEFFE-Cations were stained using a Coomassie blue fast staining solution (Beyotime, China), following general protocols.

### 4.12 Mass Spectrometry and Protein Identification

The protein components in the CEFFE-Cation group were identified by liquid chromatography-tandem mass spectrometry (LC-MS/MS) using a ChromXP Eksigent system, according to Guo *et al*. [42]. The data processing was performed with a Mascot software. We also revived technological support from the biotechnology company Oebiotech (Shanghai, China).

### 4.13 Total-Antioxidant Capacity (T-AOC) Analysis

The T-AOC of CEFFE-Cations in a cell-free system was quantified using a total antioxidant capacity assay kit (Nanjing Jiancheng Biotechnology Institute, Nanjing, China) according to the manufacturer’s protocol.

### 4.14 Intracellular ROS Measurement

Intracellular ROS levels were evaluated using a reactive oxygen species assay kit (Beyotime, China) according to the manufacturer’s instructions. Briefly, BMMs were plated and stimulated with RANKL (50 ng/mL), M-CSF (25 ng/mL), and CEFFE-Cation (100 μg/mL) for two and four days, respectively. The cells were then loaded with 10 μM DCFH-DA probe in Hank’s solution for 15 min. Images were obtained using a laser confocal fluorescence microscope (Leica TCS-SP8).

### 4.15 Immunofluorescent Staining

Nuclear translocation of NFATc1 was evaluated by immunofluorescence staining. First, BMMs were plated and stimulated with RANKL (50 ng/mL), M-CSF (25 ng/mL), and CEFFE-Cation (100 μg/mL) for two and four days, respectively. The cells were then fixed with 4% paraformaldehyde, permeabilized with 0.2% Triton, blocked with 5% BSA, and incubated with anti-NFATc1 primary antibody (1:100, Santa Cruz Biotechnology, USA) at 4°C overnight. Next, a fluorescent secondary antibody was used to visualize the NFATc1 protein, and nuclei were stained with DAPI. Images were obtained using a laser confocal fluorescence microscope (Leica TCS-SP8).

### 4.16 Intracellular Calcium Levels and Oscillation Assessment

A Fluo-4 AM probe (Beyotime, China) was used to measure intracellular Ca^2+^ levels according to the manufacturer’s protocols. Briefly, BMMs, which had been precultured for two days with the indicated treatments, were loaded with a 2 μM Fluo-4 AM probe in Hank’s solution for 30 min. After rinsing, the cells were incubated with Hank’s solution containing 5% FBS and continuously stimulated with the indicated treatments. Osteoclast precursors were excited at 488 nm, and the fluorescent signal at 510-530 was monitored at 1-sec intervals for 8 min under a Leica TCS-SP8 confocal microscope. Intracellular Ca^2+^ levels and oscillations were quantified using the Leica LAS-X software.

### 4.17 Calcineurin Activity

Cellular calcineurin activity was evaluated using a cellular calcineurin phosphatase activity assay kit (ab139464, Abcam, USA), following the manufacturer’s instructions. Briefly, BMMs were plated into 6-well plates and stimulated with RANKL (50 ng/mL), M-CSF (25 ng/mL), and CEFFE-Cation at different concentrations for one day. The cells were then lysed and desalted by gel filtration. Calcineurin substrate was added to cell extracts from pretreated cells with or without EGTA buffer. Finally, a microplate reader was used for colorimetric measurements at 620 nm.

### 4.18 Statistics

All data in this study are presented as mean ± SD. Significant differences between the two groups were analyzed using the student’s t-test. Significant differences between more than two groups were analyzed using one-way analysis of variance (ANOVA) with Tukey’s post-hoc test. Statistical significance was set at P < 0.05.

## Supporting information

Movie showing calcium oscillation in Blank group

Movie showing calcium oscillation in Ctrl group

Movie showing calcium oscillation in CEFFE-Cation group

## Supporting Information

Supporting Information is available Online or from the author.

## Acknowledgements

Yiqi Yang and Mingming Xu contributed equally to this study. This work was supported by the grants from the National Natural Science Foundation of China (Grant Nos. 11872251 and 12172223), Shanghai Municipal Science and Technology Commission (Grant No. 19140900104), Shanghai Collaborative Innovation Program on Regenerative Medicine and Stem Cell Research (Grant No. 2019CXJQ01).

## References

1. Qiu Z, Li L, Huang Y, Shi K, Zhang L, Huang C, Liang J, Zeng Q, Wang J, He X et al. Puerarin specifically disrupts osteoclast activation via blocking integrin-β3 Pyk2/Src/Cbl signaling pathway. J Orthop Translat. 2022; 33:55–69. DOI: 10.1016/j.jot.2022.01.003. PMID: 35228997.

2. Zhou F, Mei J, Yuan K, Han X, Qiao H, Tang T. Isorhamnetin attenuates osteoarthritis by inhibiting osteoclastogenesis and protecting chondrocytes through modulating reactive oxygen species homeostasis. J Cell Mol Med. 2019; 23(6):4395–4407. DOI: 10.1111/jcmm.14333. PMID: 30983153.

3. Zhao J, Yue T, Lu S, Meng H, Lin Q, Ma H, Liu G, Li H, Lu Q, Wang A et al. Local administration of zoledronic acid prevents traumatic osteonecrosis of the femoral head in rat model. J Orthop Translat. 2021; 27:132–138. DOI: 10.1016/j.jot.2020.08.005. PMID: 33786320.

4. Anam AK, Insogna K. Update on Osteoporosis Screening and Management. Med Clin North Am. 2021; 105(6):1117–1134. DOI: 10.1016/j.mcna.2021.05.016. PMID: 34688418.

5. Black DM, Rosen CJ. Clinical Practice. Postmenopausal Osteoporosis. N Engl J Med. 2016; 374(3):254–262. DOI: 10.1056/NEJMcp1513724. PMID: 26789873.

6. Wang X, Yamauchi K, Mitsunaga T. A review on osteoclast diseases and osteoclastogenesis inhibitors recently developed from natural resources. Fitoterapia. 2020; 142:104482. DOI: 10.1016/j.fitote.2020.104482. PMID: 31954740.

7. Udagawa N, Koide M, Nakamura M, Nakamichi Y, Yamashita T, Uehara S, Kobayashi Y, Furuya Y, Yasuda H, Fukuda C et al. Osteoclast differentiation by RANKL and OPG signaling pathways. J Bone Miner Metab. 2021; 39(1):19–26. DOI: 10.1007/s00774-020-01162-6. PMID: 33079279.

8. Guo W, Li H, Lou Y, Zhang Y, Wang J, Qian M, Wei H, Xiao J, Xu Y. Tyloxapol inhibits RANKL-stimulated osteoclastogenesis and ovariectomized-induced bone loss by restraining NF-κB and MAPK activation. J Orthop Translat. 2021; 28:148–158. DOI: 10.1016/j.jot.2021.01.005. PMID: 33981577.

9. Okada H, Okabe K, Tanaka S. Finely-Tuned Calcium Oscillations in Osteoclast Differentiation and Bone Resorption. Int J Mol Sci. 2020; 22(1). DOI: 10.3390/ijms22010180. PMID: 33375370.

10. Liu W, Le CC, Wang D, Ran D, Wang Y, Zhao H, Gu J, Zou H, Yuan Y, Bian J et al. Ca(2+)/CaM/CaMK signaling is involved in cadmium-induced osteoclast differentiation. Toxicology. 2020; 441:152520. DOI: 10.1016/j.tox.2020.152520. PMID: 32522522.

11. Madreiter-Sokolowski CT, Thomas C, Ristow M. Interrelation between ROS and Ca(2+) in aging and age-related diseases. Redox Biol. 2020; 36:101678. DOI: 10.1016/j.redox.2020.101678. PMID: 32810740.

12. Rendina-Ruedy E, Rosen CJ. Bone-Fat Interaction. Endocrinol Metab Clin North Am. 2017; 46(1):41–50. DOI: 10.1016/j.ecl.2016.09.004. PMID: 28131135.

13. Xu M, Du J, Cui J, Zhang S, Zhang S, Deng M, Zhang W, Li H, Yu Z. Cell-Free Fat Extract Prevents Tail Suspension-Induced Bone Loss by Inhibiting Osteocyte Apoptosis. Front Bioeng Biotechnol. 2022; 10:818572. DOI: 10.3389/fbioe.2022.818572. PMID: 35174144.

14. Yin M, Wang X, Yu Z, Wang Y, Wang X, Deng M, Zhao D, Ji S, Jia N, Zhang W. γ-PGA hydrogel loaded with cell-free fat extract promotes the healing of diabetic wounds. J Mater Chem B. 2020; 8(36):8395–8404. DOI: 10.1039/d0tb01190h. PMID: 32966542.

15. Wang X, Deng M, Yu Z, Cai Y, Liu W, Zhou G, Wang X, Cao Y, Li W, Zhang W. Cell-free fat extract accelerates diabetic wound healing in db/db mice. Am J Transl Res. 2020; 12(8):4216–4227. DOI: PMID: 32913499.

16. Xu Y, Deng M, Cai Y, Zheng H, Wang X, Yu Z, Zhang W, Li W. Cell-Free Fat Extract Increases Dermal Thickness by Enhancing Angiogenesis and Extracellular Matrix Production in Nude Mice. Aesthet Surg J. 2020; 40(8):904–913. DOI: 10.1093/asj/sjz306. PMID: 31679030.

17. Deng M, Wang X, Yu Z, Cai Y, Liu W, Zhou G, Wang X, Yu Z, Li W, Zhang WJ. Cell-free fat extract promotes tissue regeneration in a tissue expansion model. Stem Cell Res Ther. 2020; 11(1):50. DOI: 10.1186/s13287-020-1564-7. PMID: 32019588.

18. Yu Z, Cai Y, Deng M, Li D, Wang X, Zheng H, Xu Y, Li W, Zhang W. Fat extract promotes angiogenesis in a murine model of limb ischemia: a novel cell-free therapeutic strategy. Stem Cell Res Ther. 2018; 9(1):294. DOI: 10.1186/s13287-018-1014-y. PMID: 30409190.

19. Lee KM, Park KH, Hwang JS, Lee M, Yoon DS, Ryu HA, Jung HS, Park KW, Kim J, Park SW et al. Inhibition of STAT5A promotes osteogenesis by DLX5 regulation. Cell Death Dis. 2018; 9(11):1136. DOI: 10.1038/s41419-018-1184-7. PMID: 30429452.

20. Yuan K, Mei J, Shao D, Zhou F, Qiao H, Liang Y, Li K, Tang T. Cerium Oxide Nanoparticles Regulate Osteoclast Differentiation Bidirectionally by Modulating the Cellular Production of Reactive Oxygen Species. Int J Nanomedicine. 2020; 15:6355–6372. DOI: 10.2147/ijn.S257741. PMID: 32922006.

21. Yang Y, Chung MR, Zhou S, Gong X, Xu H, Hong Y, Jin A, Huang X, Zou W, Dai Q et al. STAT3 controls osteoclast differentiation and bone homeostasis by regulating NFATc1 transcription. J Biol Chem. 2019; 294(42):15395–15407. DOI: 10.1074/jbc.RA119.010139. PMID: 31462535.

22. Zeng XZ, He LG, Wang S, Wang K, Zhang YY, Tao L, Li XJ, Liu SW. Aconine inhibits RANKL-induced osteoclast differentiation in RAW264.7 cells by suppressing NF-κB and NFATc1 activation and DC-STAMP expression. Acta Pharmacol Sin. 2016; 37(2):255–263. DOI: 10.1038/aps.2015.85. PMID: 26592521.

23. Xin Y, Liu Y, Liu D, Li J, Zhang C, Wang Y, Zheng S. New Function of RUNX2 in Regulating Osteoclast Differentiation via the AKT/NFATc1/CTSK Axis. Calcif Tissue Int. 2020; 106(5):553–566. DOI: 10.1007/s00223-020-00666-7. PMID: 32008052.

24. Xian Y, Su Y, Liang J, Long F, Feng X, Xiao Y, Lian H, Xu J, Zhao J, Liu Q et al. Oroxylin A reduces osteoclast formation and bone resorption via suppressing RANKL-induced ROS and NFATc1 activation. Biochem Pharmacol. 2021; 193:114761. DOI: 10.1016/j.bcp.2021.114761. PMID: 34492273.

25. Hasegawa H, Kido S, Tomomura M, Fujimoto K, Ohi M, Kiyomura M, Kanegae H, Inaba A, Sakagami H, Tomomura A. Serum calcium-decreasing factor, caldecrin, inhibits osteoclast differentiation by suppression of NFATc1 activity. J Biol Chem. 2010; 285(33):25448–25457. DOI: 10.1074/jbc.M109.068742. PMID: 20547767.

26. Zhang J, Liu Y, Chen Y, Yuan L, Liu H, Wang J, Liu Q, Zhang Y. Adipose-Derived Stem Cells: Current Applications and Future Directions in the Regeneration of Multiple Tissues. Stem Cells Int. 2020; 2020:8810813. DOI: 10.1155/2020/8810813. PMID: 33488736.

27. Zhang D, Ni N, Wang Y, Tang Z, Gao H, Ju Y, Sun N, He X, Gu P, Fan X. CircRNA-vgll3 promotes osteogenic differentiation of adipose-derived mesenchymal stem cells via modulating miRNA-dependent integrin α5 expression. Cell Death Differ. 2021; 28(1):283–302. DOI: 10.1038/s41418-020-0600-6. PMID: 32814879.

28. Yang Y, Lin Y, Wang M, Yuan K, Wang Q, Mu P, Du J, Yu Z, Yang S, Huang K et al. Targeting ferroptosis suppresses osteocyte glucolipotoxicity and alleviates diabetic osteoporosis. Bone Res. 2022; 10(1):26. DOI: 10.1038/s41413-022-00198-w. PMID: 35260560.

29. Kajimura D, Lee HW, Riley KJ, Arteaga-Solis E, Ferron M, Zhou B, Clarke CJ, Hannun YA, DePinho RA, Guo XE et al. Adiponectin regulates bone mass via opposite central and peripheral mechanisms through FoxO1. Cell Metab. 2013; 17(6):901–915. DOI: 10.1016/j.cmet.2013.04.009. PMID: 23684624.

30. Upadhyay J, Farr OM, Mantzoros CS. The role of leptin in regulating bone metabolism. Metabolism. 2015; 64(1):105–113. DOI: 10.1016/j.metabol.2014.10.021. PMID: 25497343.

31. Kirk B, Feehan J, Lombardi G, Duque G. Muscle, Bone, and Fat Crosstalk: the Biological Role of Myokines, Osteokines, and Adipokines. Curr Osteoporos Rep. 2020; 18(4):388–400. DOI: 10.1007/s11914-020-00599-y. PMID: 32529456.

32. Stivers KB, Beare JE, Chilton PM, Williams SK, Kaufman CL, Hoying JB. Adipose-derived cellular therapies in solid organ and vascularized-composite allotransplantation. Curr Opin Organ Transplant. 2017; 22(5):490–498. DOI: 10.1097/mot.0000000000000452. PMID: 28873074.

33. Li Y, Shi Z, Jules J, Chen S, Kesterson RA, Zhao D, Zhang P, Feng X. Specific RANK Cytoplasmic Motifs Drive Osteoclastogenesis. J Bone Miner Res. 2019; 34(10):1938–1951. DOI: 10.1002/jbmr.3810. PMID: 31173390.

34. Lee JY, Kim DA, Kim EY, Chang EJ, Park SJ, Kim BJ. Lumican Inhibits Osteoclastogenesis and Bone Resorption by Suppressing Akt Activity. Int J Mol Sci. 2021; 22(9). DOI: 10.3390/ijms22094717. PMID: 33946862.

35. Kim HJ, Kim HJ, Bae MK, Kim YD. Suppression of Osteoclastogenesis by Melatonin: A Melatonin Receptor-Independent Action. Int J Mol Sci. 2017; 18(6). DOI: 10.3390/ijms18061142. PMID: 28587149.

36. Kajiya H. Calcium signaling in osteoclast differentiation and bone resorption. Adv Exp Med Biol. 2012; 740:917–932. DOI: 10.1007/978-94-007-2888-2_41. PMID: 22453976.

37. Asagiri M, Sato K, Usami T, Ochi S, Nishina H, Yoshida H, Morita I, Wagner EF, Mak TW, Serfling E et al. Autoamplification of NFATc1 expression determines its essential role in bone homeostasis. J Exp Med. 2005; 202(9):1261–1269. DOI: 10.1084/jem.20051150. PMID: 16275763.

38. Dumbuya H, Hafez SY, Oancea E. Cross talk between calcium and ROS regulate the UVA-induced melanin response in human melanocytes. Faseb j. 2020; 34(9):11605–11623. DOI: 10.1096/fj.201903024R. PMID: 32658369.

39. Zhuang X, Hu G. In vitro Osteoclastogenesis Assays Using Primary Mouse Bone Marrow Cells. Bio Protoc. 2018; 8(11):e2875. DOI: 10.21769/BioProtoc.2875. PMID: 34285989.

40. Yang Y, Wang M, Yang S, Lin Y, Zhou Q, Li H, Tang T. Bioprinting of an osteocyte network for biomimetic mineralization. Biofabrication. 2020; 12(4):045013. DOI: 10.1088/1758-5090/aba1d0. PMID: 32610301.

41. He Z, Li H, Han X, Zhou F, Du J, Yang Y, Xu Q, Zhang S, Zhang S, Zhao N et al. Irisin inhibits osteocyte apoptosis by activating the Erk signaling pathway in vitro and attenuates ALCT-induced osteoarthritis in mice. Bone. 2020; 141:115573. DOI: 10.1016/j.bone.2020.115573. PMID: 32768686.

42. Guo Z, Dai Y, Hu W, Zhang Y, Cao Z, Pei W, Liu N, Nie J, Wu A, Mao W et al. The long noncoding RNA CRYBG3 induces aneuploidy by interfering with spindle assembly checkpoint via direct binding with Bub3. Oncogene. 2021; 40(10):1821–1835. DOI: 10.1038/s41388-020-01601-8. PMID: 33564066.

