## Supporting Information for "Fat Extract Modulates Calcium Signaling and Protects Against Hyperactive Osteoclastogenesis in Bone Remodeling with Antioxidant Capacity"

**Figure S1.** Effect of CEFFE on BMMs viability

**Figure S2.** CEFFE does not inhibit RANKL-induced ERK/AKT phosphorylation

**Figure S3.** CEFFE does not inhibit RANKL-induced NF-κB activation

**Table S1.** Primer sequences used in RT-qPCR

**Legend of Movie S1.** Movie showing calcium oscillation signals in Blank group

**Legend of Movie S2.** Movie showing calcium oscillation signals in Ctrl group

**Legend of Movie S3.** Movie showing calcium oscillation signals in CEFFE-Cation group

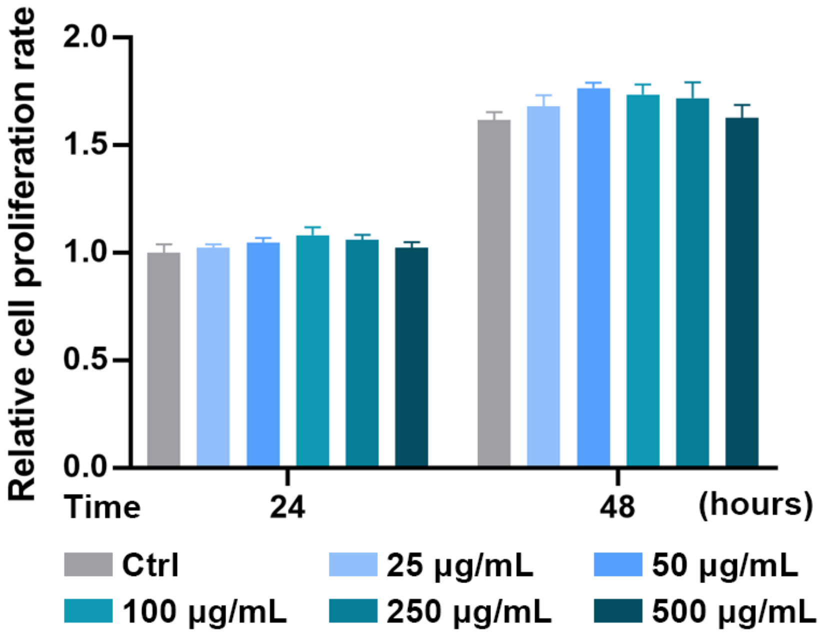


**Figure S1.** Effect of CEFFE on BMMs viability. BMMs treated with various concentrations of CEFFE for 24 h and 48 h were assessed by CCK-8 assay.

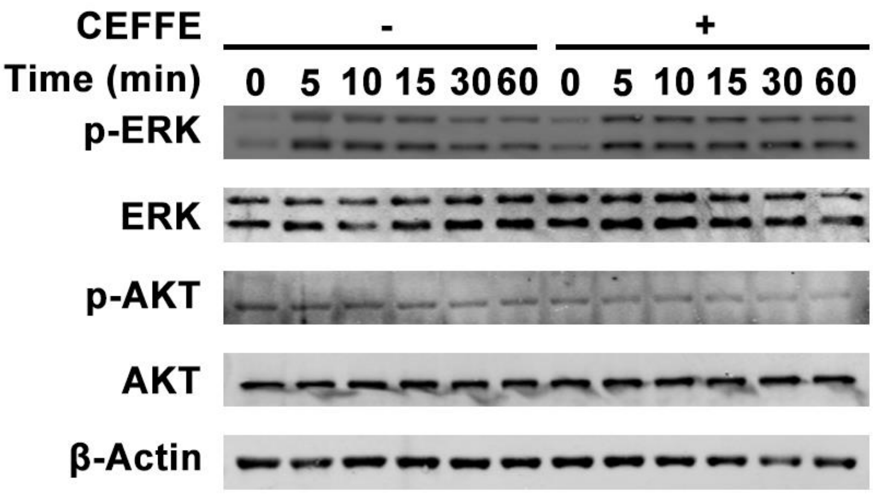


**Figure S2.** CEFFE does not inhibit RANKL-induced ERK/AKT phosphorylation. Western blot analysis of ERK, p-ERK, AKT, p-AKT. BMMs pretreated with/without CEFFE were stimulated with RANKL for 60 min. No obvious inhibition effect was detected.

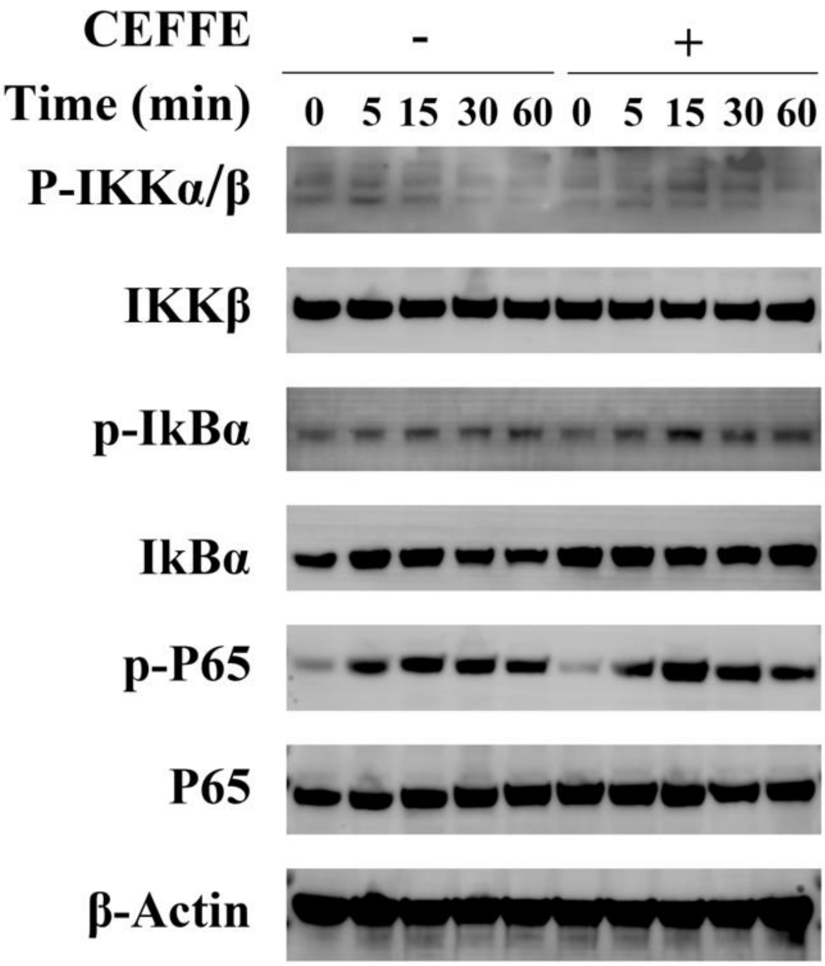


**Figure S3.** CEFFE does not inhibit RANKL-induced NF-κB activation. Western blot analysis of the NF-κB pathway. BMMs pretreated with/without CEFFE were stimulated with RANKL for 60 min. No obvious inhibition effect was detected.

**Table S1.** Primer sequences used in RT-qPCR

| Gene | Sequence | |
|---|---|---|
| β-Actin | Forward | AGAGGGAAATCGTGCGTGACA |
| | Reverse | CACTGTGTTGGCATAGAGGTC |
| Trap | Forward | CGCTGCCTTGTCAAGAACTT |
| | Reverse | CGTTGATGTCGCACAGAGG |
| Ctsk | Forward | GGACCCATCTCTGTGTCCAT |
| | Reverse | CCGAGCCAAGAGAGCATATC |
| Dcstamp | Forward | AAAACCCTTGGGCTGTTCTT |
| | Reverse | AATCATGGACGACTCCTTGG |
| Nfatc1 | Forward | GGGTCAGTGTGACCGAAGAT |
| | Reverse | GGAAGTCAGAAGTGGGTGGA |
| Atp6a3 | Forward | CACAGGGTCTGCTTACAACTG |
| | Reverse | CGTCTACCACGAAGCGTCTC |
| Atp6d2 | Forward | TGCGGCAGGCTCTATCCAGAGG |
| | Reverse | CCACTGCCACCGACAGCGTC |
| Calm1 | Forward | TGGGAATGGTTACATCAGTGC |
| | Reverse | CGCCATCAATATCTGCTTCTCT |
| Rcan1 | Forward | TTGTGTGGCAAACGATGATGT |
| | Reverse | CCCAGGAACTCGGTCTTGT |
| Calna | Forward | ATCCCAAGTTGTCGACGACC |
| | Reverse | ACACTTTCTTCCAGCCTGCC |
| Cnb1 | Forward | TGCCTGAGTTACAGCAGAACC |
| | Reverse | TCGCCTTTGACACTGAACTG |

**Movie S1.** Movie showing calcium oscillation signals in Blank group. BMMs pretreated with PBS were loaded with Fluo-4AM probe. No significant $Ca^{2+}$ oscillation signals were detected.

**Movie S2.** Movie showing calcium oscillation signals in Ctrl group. BMMs pretreated with RANKL (50 ng/mL) were loaded with Fluo-4AM probe. Obvious RANKL-evoked $Ca^{2+}$ oscillation signals were detected.

**Movie S3.** Movie showing calcium oscillation signals in CEFFE-Cation group. BMMs pretreated with RANKL (50 ng/mL) and CEFFE-Cation (100 μg/mL) were loaded with Fluo-4AM probe. CEFFE-Cation efficiently inhibited RANKL-evoked $Ca^{2+}$ oscillation signals.
